# METTL1 regulates glioma proliferation through internal m7G methylation of EPHA2

**DOI:** 10.64898/2026.07.05.736545

**Authors:** Tutu Xu, Peng Yu, Yongqing Sun, Jinhai Huang, Xiang Fang, Junhui Lv, Shuxu Yang, Guangyu Li

**Affiliations:** Department of Neurosurgery, the First Affiliated Hospital of China Medical University, No. 155, North Nanjing Street, Heping District, Shenyang, Liaoning 110001, China; Department of Neurosurgery, Sir Run Run Shaw Hospital, School of Medicine, Zhejiang University, Hangzhou, Zhejiang, China; Department of Neurosurgery, Central hospital affiliated to Shandong First Medical University, Jinan, Shandong, China

**Keywords:** METTL1, EPHA2, m7G modification, mRNA stability, AKT signaling pathway, Glioma proliferation

## Abstract

**Background:** Methyltransferase-like 1 (METTL1) is highly expressed in organs like the pancreas but less so in the brain. The METTL1-WDR4 complex catalyzes N7-methylguanosine (m7G) methylation in tRNA, miRNA, mRNA, and rRNA, which impacts RNA stability and function. These modifications affect mRNA translation and tRNA functionality, influencing protein production and cellular activities. Such modifications can regulate tumor growth, invasion, and metabolism by selectively controlling protein expression.

**Method:** Gene expression data from public databases were analyzed to compare METTL1 expression in normal and tumor tissues. Western blot (WB) and immunohistochemistry (IHC) were used to quantify METTL1 levels in glioma samples and assess their prognostic significance. Cell viability, migration, invasion, and proliferation were evaluated using Cell Counting Kit-8 (CCK-8), wound healing, Transwell, cell cycle analysis, and colony formation assays. RNA immunoprecipitation PCR (RIP-PCR) identified m7G methylation sites on EPHA2 mRNA, and RNA stability was assessed with actinomycin D.

**Results:** Bioinformatics analysis revealed that METTL1 is overexpressed in gliomas, correlating with poor prognosis. Knockdown of METTL1 significantly affected cell proliferation, migration, and invasion. RNA sequencing (RNA-seq) and m7G analysis identified EPHA2 as a downstream target, influencing the cell cycle via the AKT pathway. RIP and methylated RNA immunoprecipitation (MeRIP) confirmed two m7G sites on EPHA2 mRNA regulated by METTL1. Small interfering RNA (siRNA)-mediated METTL1 knockdown in EPHA2 mutants affected mRNA stability. Rescue experiments restored cell proliferation and AKT pathway gene expression.

**Conclusion:** METTL1 methylates EPHA2 mRNA, enhancing its stability and expression, which activates the AKT signaling pathway and influences glioma cell proliferation. METTL1 could be a potential therapeutic target in glioma treatment.

## Introduction

Gliomas are among the most common primary brain tumors in adults, accounting for nearly 30% of all primary brain tumors and 80% of malignant ones, making them the leading cause of death from primary brain tumors(1,2). Glioblastoma multiforme (GBM) is the most prevalent and malignant form (WHO grade IV) with an incidence of 3.2 per 100,000 people(3). Despite standardized treatments, patient prognosis remains poor(3–5). Thus, understanding the molecular mechanisms of gliomas and finding effective therapeutic targets are critical.

Recent studies have highlighted the crucial role of epigenetic modifications in glioma development(6–8), particularly methylation abnormalities, which are linked to different glioma subtypes. For instance, IDH mutations are strongly associated with the cytosine-phosphate-guanine (CpG) island methylator phenotype (G-CIMP)(9–11), common in low-grade gliomas and associated with better prognosis(12,13). Methylation of the O-6-Methylguanine-DNA Methyltransferase (MGMT) gene promoter is a favorable prognostic factor in GBM, especially with alkylating agent chemotherapy like temozolomide(12–14).

As tumor epigenetics research advances, RNA methylation, beyond DNA methylation, has gained focus. RNA methylation is a well-known RNA modification, with over 70 types identified, including N6-methyladenosine (m6A), N1-methyladenosine (m1A), N5-methylcytosine (m5C), and N7-methylguanosine (m7G)(15–17). While m6A is the most extensively studied, m7G has attracted increasing attention recently. m7G is a ubiquitous and evolutionarily conserved RNA modification, first found in the 5’ cap of mRNA(18,19). Most eukaryotic mRNAs have an m7G cap at the 5’ end(17,20,21), which stabilizes the transcript, prevents degradation(22–24), and mediates various functions like transcription elongation(25), pre-mRNA splicing(26,27), polyadenylation(28), nuclear export(29), and translation(30).

Beyond the mRNA cap, m7G is also found internally in mRNA, tRNA, and rRNA(31,32). Internal m7G modifications play critical roles in RNA metabolism, including processing, stability, maturation, and translation(33,34). These modifications are dynamically regulated by methyltransferases, with METTL1 being a key enzyme. METTL1, the mammalian homolog of yeast Trm8, forms a functional methyltransferase complex with WDR4, the human homolog of yeast Trm82, to mediate m7G methylation(32).

Despite significant progress in m7G research, much about internal m7G modifications’ locations and functions remains unknown. Internal m7G is associated with disease-related protein dysregulation, suggesting its role as an epitranscriptomic marker that could lead to various neurological diseases(35–38). This study combines bioinformatics analysis and clinical data to reveal the close association between METTL1 and gliomas, indicating its potential involvement in glioma proliferation, invasion, and metastasis. Further experiments explore METTL1’s role in internal m7G methylation mechanisms.

## Materials and methods

### Bioinformatic analysis

RNA-seq and clinical data for glioma patients were obtained from TCGA and normal brain tissue data from GTEx via UCSC Xena (http://xena.ucsc.edu). Additional mRNA expression and clinical data for glioma patients were sourced from the CGGA (http://www.cgga.org.cn). In R Studio (Boston, MA, USA), PRMT6 and YTHDF2 expression and survival analyses were performed. Differential gene expression was analyzed using the Limma R package, and Gene Set Enrichment Analysis (GSEA) was conducted with the clusterProfiler package.

### Cell lines and cell culture conditions

Human GBM cell lines (LN229, U251MG, U87MG, U118MG), the normal brain glial cell line (HEB), and HEK293T cells were sourced from the Cell Bank of the Chinese Academy of Sciences (Shanghai, China). U87MG cells were cultured in Eagle’s Minimum Essential Medium (EMEM) with 10% fetal bovine serum (FBS) at 37°C and 5% CO₂. All other cell lines were maintained in Dulbecco’s Modified Eagle Medium (DMEM) with 10% FBS under identical conditions.

### Lentivirus packaging and infection

Lentiviral shRNA plasmids targeting METTL1, EPHA2, PTTG1 and PPIC, along with plasmids expressing full-length METTL1 and EPHA2, were constructed and transfected into HEK-293T cells using Lipofectamine 8000 (Beyotime, Cat#C0533) following the manufacturer’s protocol. The plasmid ratio for transfection was pVSVG:psPAX2:target plasmid = 1:3:4. Virus-containing supernatants were collected at 36 and 48 hours post-transfection and used to infect target cells in the presence of 10 µg/mL polybrene (Solarbio).

Infected cells were selected with 2 µg/mL puromycin for three days to isolate successfully transduced cells. For rescue experiments, double-stable cell lines were established by combining standard puromycin selection (2 µg/mL for three days) with 800 µg/mL G418 selection for seven days.

### Western blot

Cells were lysed with RIPA lysis buffer containing protease and phosphatase inhibitors (Beyotime, Cat# P0013B) according to the manufacturer’s instructions. Protein concentrations were determined using the BCA Protein Assay Kit (Beyotime, Cat# P0010). Equal protein amounts were resolved by SDS-PAGE (Tris-HCl) and transferred onto PVDF membranes. Membranes were blocked with 5% skim milk at room temperature for 1 hour, then incubated overnight at 4°C with primary antibodies, followed by HRP-conjugated secondary antibodies for 1 hour at room temperature. Chemiluminescent signals were detected using Enhanced Chemiluminescence (ECL) and quantified with ImageJ software (NIH, Bethesda, USA).

### Quantitative real-time PCR assay (qPCR)

Total RNA was extracted from cells using RNAiso Plus reagent (TaKaRa, Cat# 9109) following the manufacturer’s protocol. cDNA was synthesized from RNA using the abm ALL-In-One 5X RT MasterMix kit (abm, Cat# G592). Quantitative PCR (qPCR) was performed using the BlasTaq™ 2X qPCR MasterMix kit (abm, Cat# G891), with β-actin as the internal reference. RNA expression levels were calculated using the 2^−ΔΔCT^ method. Primer sequences were synthesized by BGI (Beijing, China) and are provided in Supplementary Material 4: Table S1.

### RNA immunoprecipitation (RIP)assays

The RIP experiment was performed using the EZ-Magna RIP RNA-Binding Protein Immunoprecipitation Kit (Millipore, Cat#17–701) according to the manufacturer’s protocol. Cells were collected with a scraper and lysed in RIP lysis buffer. Protein A/G magnetic beads were pre-washed with RIP wash buffer and incubated with 5 µg of either specific antibody or negative control IgG at room temperature for 30 minutes. The RIP lysate was added to the antibody-bead complex (10% of the lysate was reserved as input and stored at -80°C) and incubated overnight at 4°C. The beads were collected using a magnetic stand, and the supernatant was discarded. Proteinase K buffer was used to elute the RNA from the beads, and the supernatant was transferred to a new tube for RNA extraction using RNAiso Plus. The extracted RNA was reverse transcribed to cDNA and quantified by qPCR. Primer sequences used for RIP are provided in Supplementary Material 4: Table S1.

### MeRIP-qPCR

Total RNA was extracted from cells overexpressing METTL1 or EPHA2, then fragmented to approximately 300 nt using RNA Fragmentation Reagents (Invitrogen, Cat#AM8740). The fragmented RNA was recovered using an RNA purification column (Zymo Research, Cat#R1017). To block Protein A/G magnetic beads, BSA was mixed with the beads and rotated at 4°C for 2 hours. Fifty micrograms of fragmented RNA were mixed with 5 µg of m6A antibody or IgG control and incubated with the blocked beads overnight at 4°C. After collecting the beads using a magnetic stand and discarding the supernatant, the bound RNA was eluted and digested with proteinase K. The eluted RNA was extracted with an RNA purification column, reverse transcribed, and quantitatively analyzed by qPCR. Primers used for MeRIP experiments are provided in Supplementary Material 4: Table S1.

## Results

### METTL1 is highly expressed in gliomas and is associated with poor prognosis

Analysis of METTL1 expression data from the GTEx database revealed that METTL1 expression is relatively low in normal brain tissue but significantly increased in GBM (**Fig. 1A**). This finding was consistent with analyses from the TCGA and CGGA databases, showing that METTL1 expression is higher in tumor cells compared to normal cells and increases with glioma grade (**Fig. 1B-C**). Additionally, high METTL1 expression was significantly associated with poor patient prognosis (**Fig. 1D**). To validate the consistency of bioinformatics analysis with clinical data, this study collected samples from 4 peritumor tissues, 16 low-grade glioma (LGG) tissues, and 24 GBM tissues. IHC was used to detect METTL1 expression levels in different grades of gliomas. Statistical analysis indicated a positive correlation between METTL1 expression and glioma grade in the clinical samples. Furthermore, Western Blot analysis of 4 randomly selected samples from each group confirmed that METTL1 expression levels were consistent with IHC results and bioinformatics analysis from public databases (**Fig. 1E-G**). These findings suggest that METTL1 expression levels are closely related to glioma occurrence and progression.

**Figure 1.**
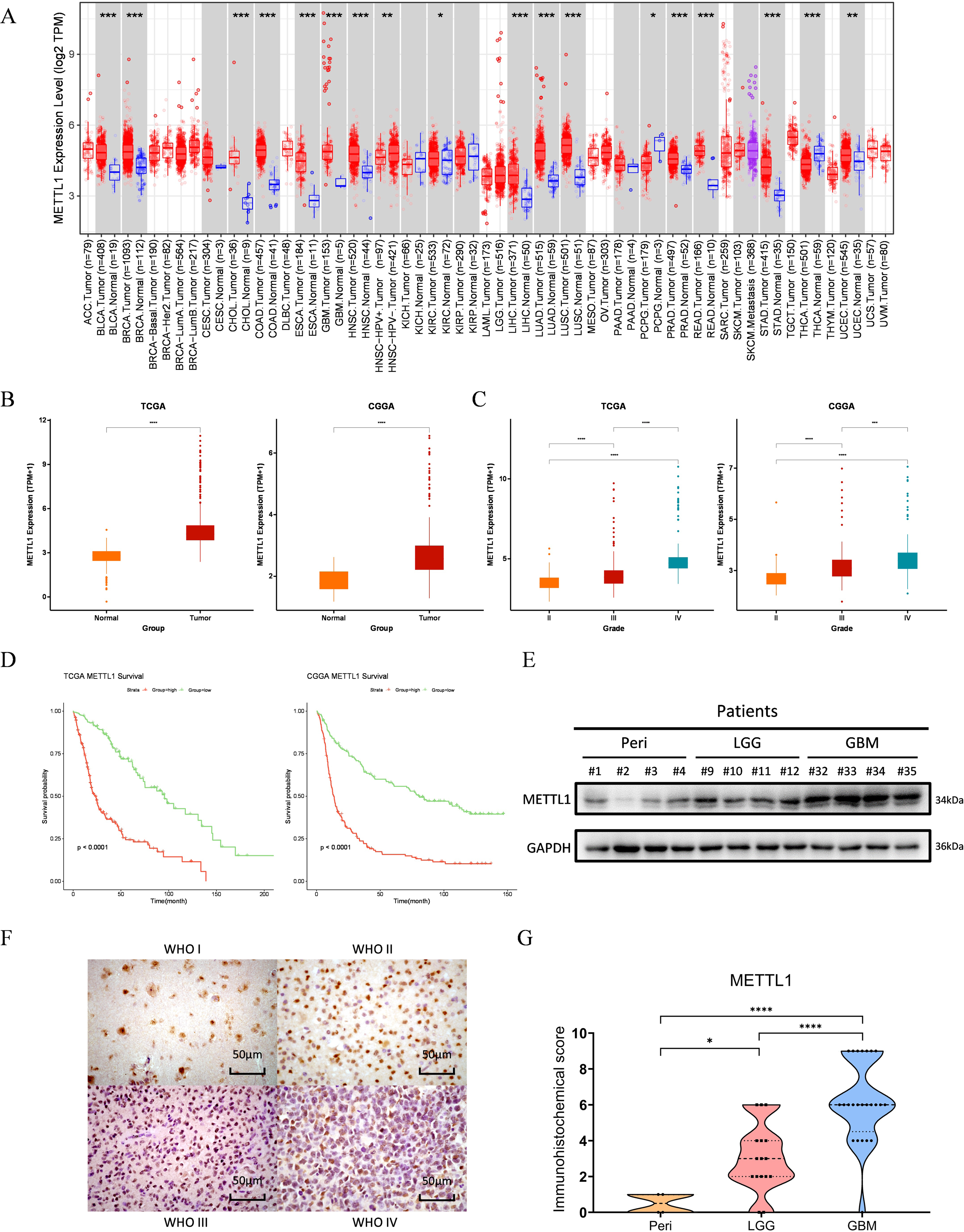
High expression of METTL1 in gliomas and its association with poor prognosis. A, Pan-cancer analysis of METTL1 expression abundance in the GTEx dataset. B and C, Differential expression of METTL1 between tumor and normal tissues, as well as across different glioma grades, in the TCGA and CGGA databases. D, Impact of different METTL1 expression levels on the survival of glioma patients in the TCGA and CGGA databases. E, Differential expression of METTL1 protein levels in clinical samples assessed by Western Blot. F, Immunohistochemical staining of METTL1 in glioma tissue samples of different grades. G, Statistical analysis of IHC scoring levels for each sample. *, P < 0.05; **, P < 0.01; ***, P < 0.001; ****, P < 0.0001

### METTL1 Knockdown Significantly Alters Glioma Cell Phenotypes

We used the widely studied glioma cell lines U251 and LN229 as in vitro models. METTL1 was knocked down and overexpressed in these cells. Using lentiviral infection, we introduced knockdown control virus (METTL1-Ctr), two different knockdown shRNA lentiviruses (sh32 and sh33), overexpression control virus (OE-NC), and overexpression virus (OE) to establish stable transfected cell lines (Supplementary Fig. S1A). Total RNA was extracted from the stable cell lines, and qPCR was performed (Supplementary Fig. S1C). Western Blot analysis confirmed that METTL1 protein levels were significantly downregulated in the knockdown groups and upregulated in the overexpression groups (**Fig. 2A**), consistent with the qPCR results. Dot Blot analysis was used to observe m7G modification levels in U251 and LN229 cells with stable METTL1 expression (**Fig. 2B**).Previous studies have identified METTL1 as an oncogene that promotes cell proliferation by regulating the cell cycle through m7G modification of tRNA(39). To investigate whether METTL1 contributes to glioma cell proliferation and malignancy, we performed CCK-8 assays on U251 and LN229 glioma cell lines with METTL1 knockdown or overexpression. Knockdown significantly inhibited cell growth in both cell lines, while overexpression accelerated proliferation only in LN229 cells, with no significant effect observed in U251 cells (**Fig. 2C**). Colony formation assays corroborated these findings (**Fig. 2D**; Supplementary Fig. S1B), demonstrating reduced clonogenic potential upon METTL1 knockdown in both cell lines.Further functional assays demonstrated that METTL1 knockdown impaired glioma cell migration and invasion (**Fig. 2E**). In wound healing assays, U251 cells with reduced METTL1 expression exhibited significantly decreased migration, whereas METTL1 overexpression did not enhance migration (Supplementary Fig. S1D). Similar trends were observed in transwell migration and invasion assays (**Fig. 2F-G**; Supplementary Fig. S1E).

**Figure 2.**
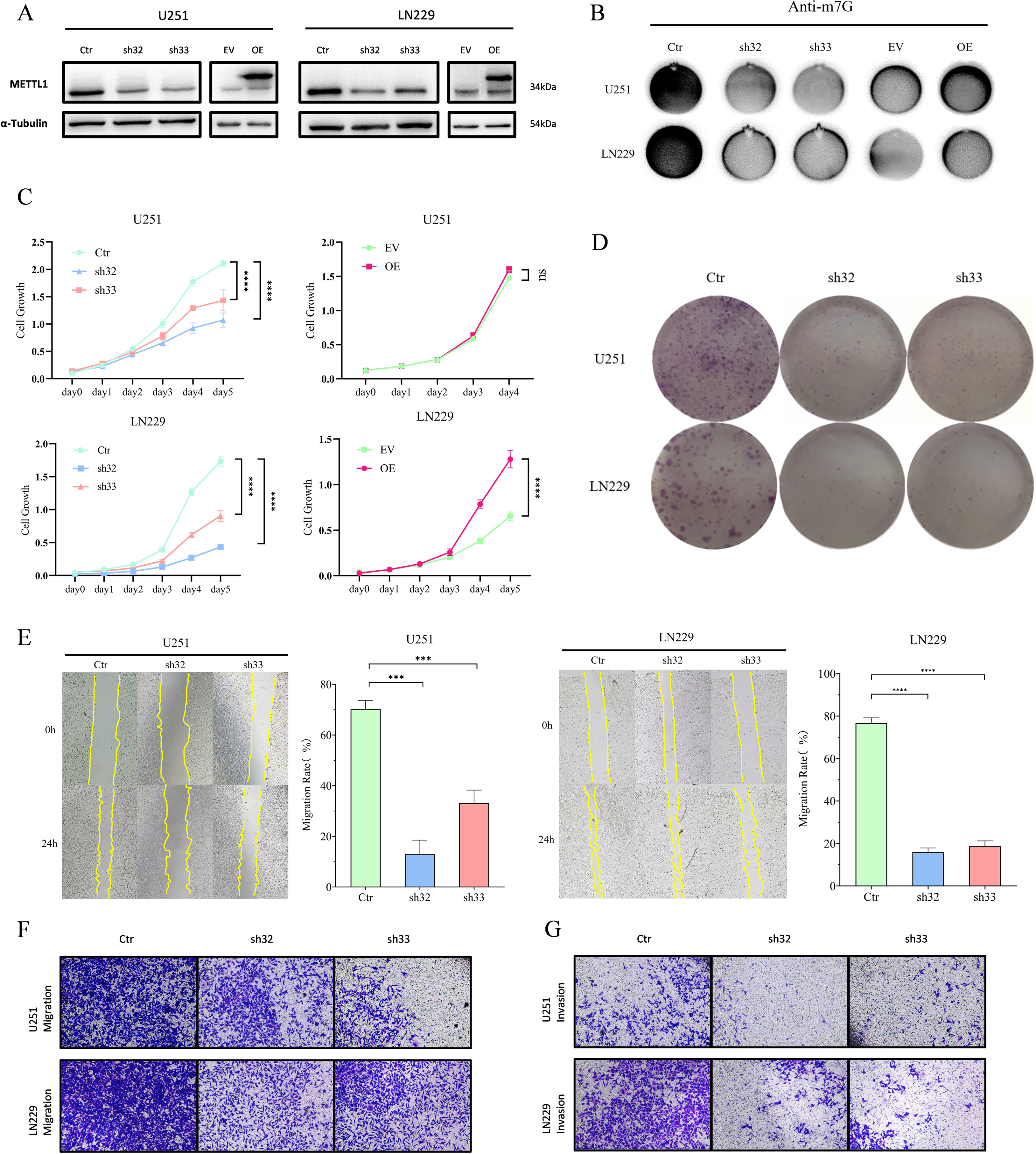
The abundance of METTL1 affects the phenotype of glioma cells. A, WB analysis of METTL1 knockdown and overexpression efficiency in cell lines. B, Dot Blot analysis was used to detect changes in m7G levels. C. CCK8 assay D. Colony formation assay. E. Scratch assay. F. Transwell assay was used to assess the migratory ability. Compared to the Ctr group, the number of transmembrane cells in the METTL1 knockdown sh32 and sh33 groups was significantly reduced. G. The invasive ability of METTL1 knockdown sh32 and sh33 cells through matrigel was significantly reduced.

### METTL1 regulates cell proliferation through multiple pathways

To explore the underlying mechanisms, we hypothesized that METTL1 might regulate glioma cell proliferation by modulating the cell cycle. Consistent with previous studies reporting G2/M phase inhibition by METTL1(40), cell cycle analysis revealed an increased proportion of cells in the G0/G1 phase following METTL1 knockdown in both U251 and LN229 cells (**Fig. 3A**), indicating G1-phase arrest. These results suggest that METTL1 may exert tumor-specific effects by targeting distinct downstream pathways.

**Figure 3.**
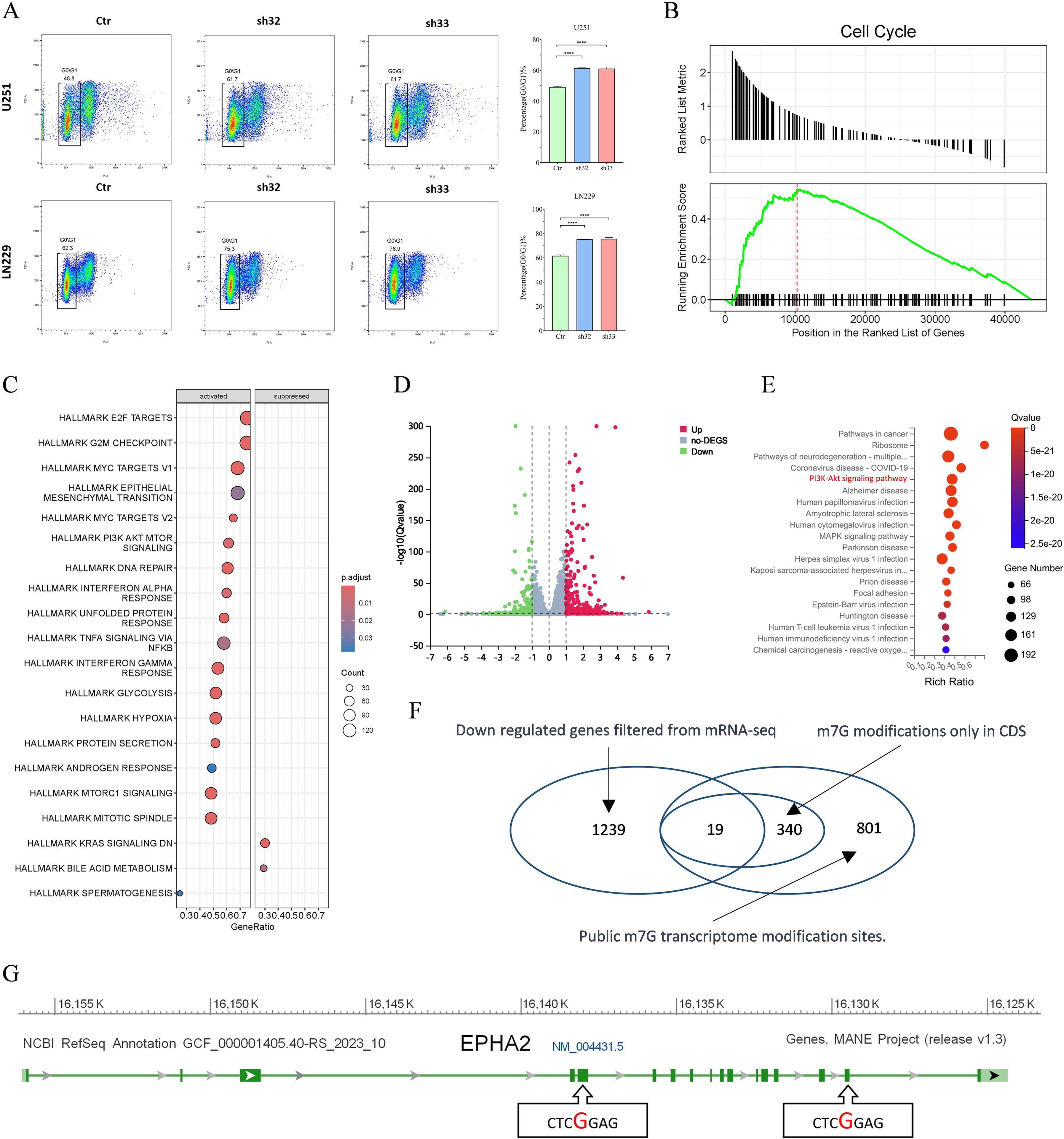
High-throughput sequencing data of METTL1 and downstream target screening. A, Cell cycle flow cytometry analysis and G0/G1 phase ratio statistics of U251 and LN229 cells following METTL1 knockdown. B, GSEA enrichment plots of METTL1 in the TCGA database. C. KEGG pathway analysis of METTL1 in the TCGA database. D. Volcano plot of differentially expressed genes following high-throughput sequencing. E. KEGG pathway analysis of differentially expressed genes from high-throughput sequencing. F. The downregulated genes obtained from sequencing were filtered and intersected with genes that have m7G modification sites exclusively in the CDS region. G. Two predicted modification sites with identical sequences in the CDS region of EPHA2.

KEGG pathway analysis using TCGA data revealed significant associations between METTL1 and multiple cell cycle-related signaling pathways (**Fig. 3B**). GSEA enrichment analysis further demonstrated that cell cycle-related genes were among the most upregulated with increased METTL1 expression (**Fig. 3C**). High-throughput sequencing of METTL1-knockdown stable cell models identified 6886 downregulated and 6315 upregulated genes (**Fig. 3D**). KEGG pathway analysis of these differentially expressed genes, combined with TCGA data, highlighted enrichment in the PI3K-AKT pathway (**Fig. 3E**). This pathway, closely linked to cell proliferation and the cell cycle, supports our observations that METTL1 promotes glioma cell proliferation.

Further analysis of sequencing data (filtering for genes with average TPM ≥0.5 in control groups and ≥30% reduction in expression following METTL1 knockdown) identified 1239 downregulated genes (**Fig. 3F**). Cross-referencing these genes with known m7G modification sites focused our attention on mRNA CDS region modifications. Screening a public m7G database yielded 340 genes with CDS-specific m7G modifications(31). Intersecting these with our downregulated genes identified 19 candidates, including EPHA2, PTTG1, and PPIC, which are highly expressed in gliomas and implicated in cell cycle regulation.Bioinformatics analysis demonstrated that elevated expression of EPHA2, PTTG1, and PPIC correlated with poor prognosis and higher glioma grades (Supplementary Fig. S2A-B). METTL1 expression showed strong RNA-level correlations with these genes (Supplementary Fig. S2C), suggesting potential regulatory interactions. Notably, EPHA2 contains two predicted m7G modification sites (CTCGGAG) in its CDS region (**Fig. 3G**; Supplementary Fig. S2D-E), underscoring its likely regulation by METTL1. EPHA2 has been extensively linked to activation of the AKT signaling pathway(41,42).

### METTL1 participates in the AKT pathway by enhancing EPHA2 mRNA Stability

We used METTL1-stable cell lines derived from U251 and LN229 glioma cells to assess the impact of METTL1 on the expression of downstream genes. qPCR analysis revealed that METTL1 knockdown significantly reduced the mRNA levels of EPHA2, PTTG1, and PPIC in both U251 and LN229 cells (**Fig. 4A**; Supplementary Fig. S3A), suggesting that METTL1 may directly or indirectly regulate these mRNAs, which contain predicted m7G modification sites. Western blot analysis confirmed that EPHA2 protein levels were significantly reduced in METTL1 knockdown cells, along with key proteins involved in the AKT signaling pathway, including p-AKT, p-mTOR, and Cyclin D1 (**Fig. 4B**). Consistent with in vitro findings, clinical glioma samples revealed that EPHA2 expression correlated positively with glioma grades (**Fig. 4C**).

**Figure 4.**
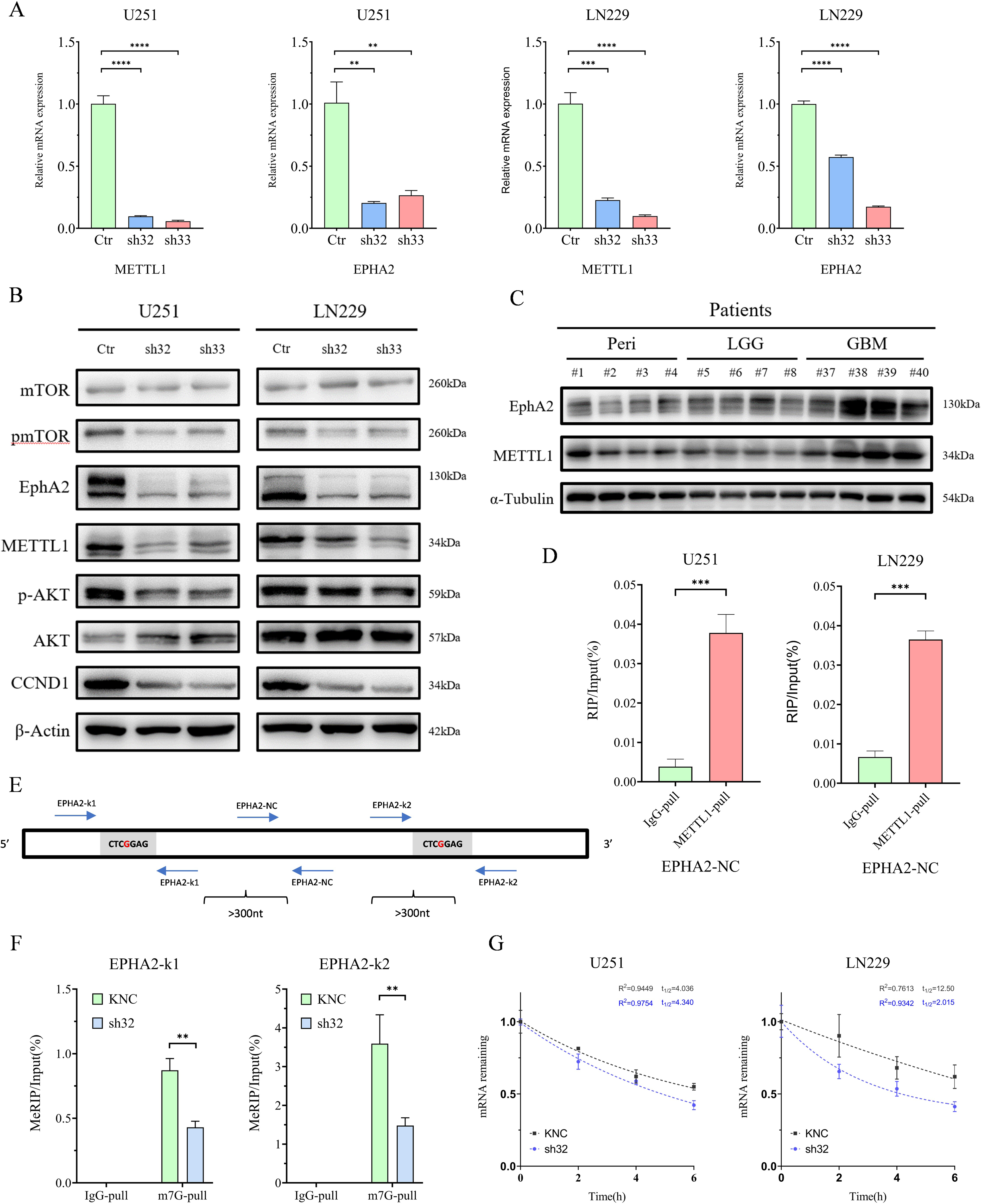
METTL1 methylates EPHA2, affecting its mRNA stability. A, Knockdown of METTL1 in U251 cells led to a decrease in EPHA2 mRNA expression levels. B, METTL1 knockdown led to changes in EPHA2, key proteins in the AKT pathway, and the downstream cell cycle protein Cyclin D1. C. Clinical sample validation showed that EPHA2 expression in gliomas increased with glioma grade progression. D. RIP assay revealed that the abundance of EPHA2 mRNA pulled down with the METTL1 antibody was significantly higher compared to IgG. E. Three primers designed for EPHA2 were spaced more than 300 nucleotides apart. F. MeRIP-PCR experiments validated the presence of m7G modification sites in two sequences between the METTL1 knockdown group and the control group. G. METTL1 knockdown led to reduced RNA stability of EPHA2.

To confirm the interaction between METTL1 and these mRNAs, RNA immunoprecipitation (RIP) was performed. RIP assays demonstrated that METTL1 protein specifically bound to the mRNAs of EPHA2, PTTG1, and PPIC, as significantly higher levels of these mRNAs were pulled down using an anti-METTL1 antibody compared to control IgG (**Fig. 4D**; Supplementary Fig. S3B). These results establish a binding interaction between METTL1 and its downstream targets.

To further validate the role of m7G methylation in regulating EPHA2 mRNA, we performed methylated RNA immunoprecipitation (MeRIP) assays. Since MeRIP requires RNA fragmentation into 300-nucleotide segments, we designed three primer pairs targeting EPHA2 (**Fig. 4E**). Primers k1 and k2 were designed to amplify regions containing predicted m7G sites, while EPHA2-NC primers targeted non-methylated control regions. qPCR analysis showed significantly reduced pull-down of EPHA2 mRNA fragments by the m7G antibody in METTL1 knockdown cells compared to controls (**Fig. 4F**). The EPHA2-NC primers amplified no specific product, confirming the absence of m7G modifications in the control region. These findings demonstrate that METTL1 mediates m7G methylation of EPHA2 mRNA at two specific sites, with the k2 site exhibiting higher modification abundance than the k1 site. This discrepancy may explain why only one m7G modification site for EPHA2 was identified in public databases.We hypothesized that METTL1 might enhance mRNA stability through m7G methylation. Actinomycin D treatment of METTL1 knockdown and control cells confirmed this hypothesis, as EPHA2 mRNA stability was significantly reduced in METTL1 knockdown cells (**Fig. 4G**).

Given that the m7G modification sites of PTTG1 and PPIC are located at shared G bases within the coding sequence, making mutations challenging to interpret, we focused further mechanistic studies on EPHA2. This gene is well-characterized for its role in the AKT signaling pathway and contains synonymous G base sites that facilitate mutational studies.

### Single-base validation of the m7G modification site in EPHA2

While METTL1’s role in tRNA methylation and its impact on translational regulation is well-documented, direct evidence linking m7G methylation to EPHA2 mRNA stability remained lacking. To address this, we constructed lentiviral plasmids encoding wild-type and mutant EPHA2 transcripts, including wild-type (E2-Wt), single-site mutants (E2-Mut1 and E2-Mut2), and a double-site mutant (E2-Mut1+2), where predicted m7G sites (G bases) were synonymously mutated to C bases to avoid nonsynonymous mutation (**Fig. 5A**). A 3xFlag tag was added at the 3′ end of the EPHA2 transcript for specific detection (**Fig. 5E**).

**Figure 5.**
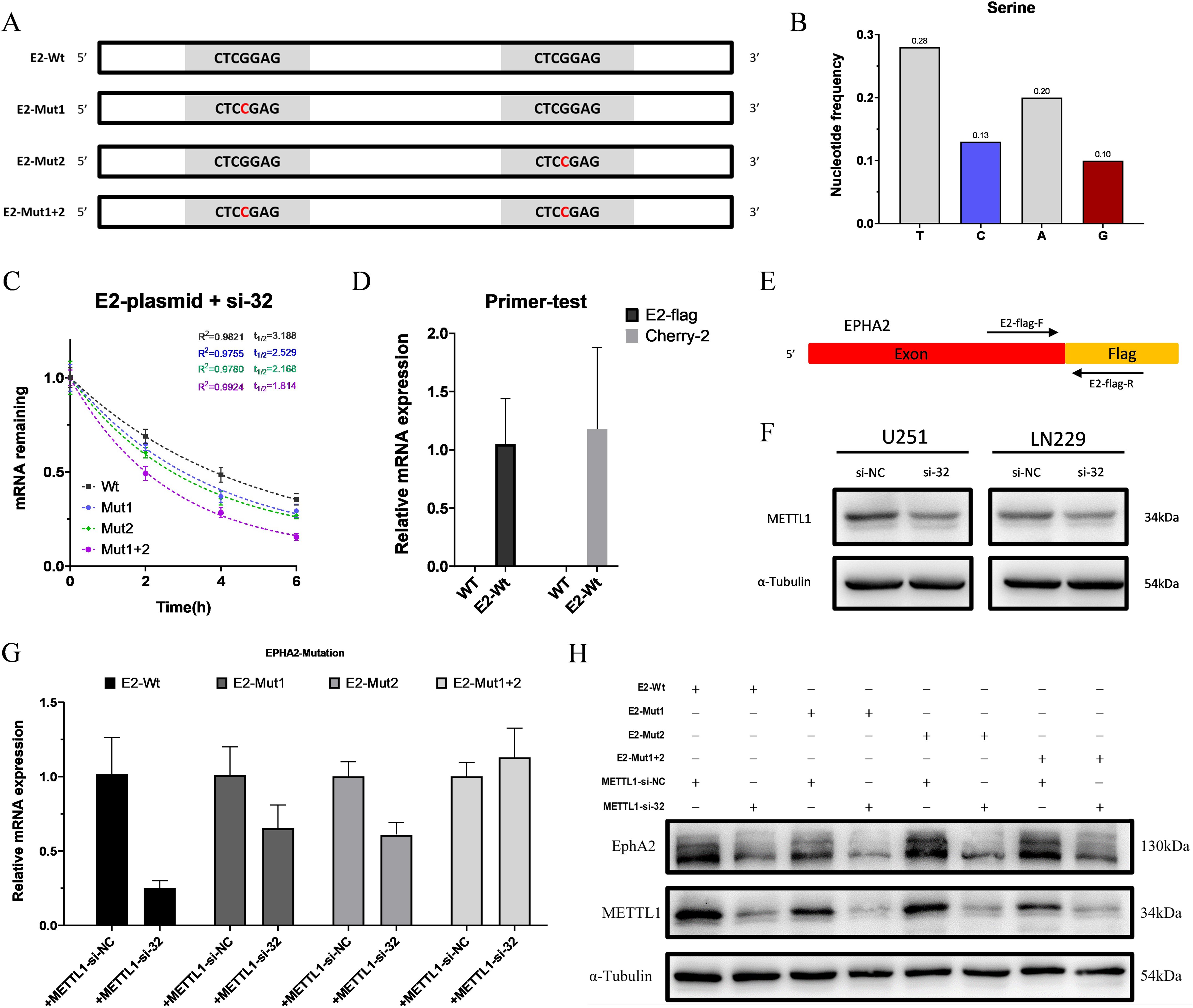
Introduction of exogenous EPHA2 and its mutants, followed by functional validation. A, EPHA2 wild-type and mutant forms. B, Selection of the base mutations for the mutant form. C. RNA stability assay of exogenous EPHA2 following METTL1 knockdown. D. RIP assay revealed that the abundance of EPHA2 mRNA pulled down with the METTL1 antibody was significantly higher compared to IgG. E. Specific detection methods for exogenous EPHA2. F. METTL1 knockdown validation using si-32, which shares the same sequence as sh32.. G. qPCR was used to verify the changes in mRNA expression levels of exogenous EPHA2 following METTL1 knockdown. H. Western Blot was used to verify the changes in protein expression levels in each group before and after METTL1 knockdown.

During lentiviral packaging and infection, variability in exogenous EPHA2 expression was observed despite controlled viral titers and infection conditions. To account for this, we normalized EPHA2 expression using the independently expressed mCherry gene as an exogenous reference (Supplementary Fig. S3C). Verification by qPCR confirmed that neither mCherry nor E2-flag transcripts were detected in the U251 cells (**Fig. 5D**), ensuring specific detection of exogenous EPHA2.We then knocked down METTL1 in each stable cell line expressing exogenous wild-type and mutant EPHA2 using validated siRNA si-32 (**Fig. 5F**) and observed the expression changes. qPCR results showed that compared to E2-Wt, the mRNA expression of E2-Mut1+2, which lacks both methylatable G bases, was almost unaffected by METTL1 knockdown, while E2-Mut1 and E2-Mut2 single-point mutants had intermediate levels of mRNA reduction (**Fig. 5G**). RNA stability assays post-METTL1 knockdown mirrored these findings: E2-Mut1+2 showed the lowest RNA stability due to the absence of m7G modification, followed by the single-site mutants, with E2-Wt exhibiting the highest stability (**Fig. 5C**). Finally, we assessed the protein expression levels of EPHA2 and its mutants before and after METTL1 knockdown (**Fig. 5H**). The Western Blot analysis confirmed METTL1 knockdown but was not used for mRNA translation regulation analysis due to METTL1’s known tRNA methylation function.

### EPHA2 Rescues METTL1 Knockdown-Induced Impaired Glioma Cell Proliferation

In the stable METTL1 knockdown model sh32 in U251 cells, we introduced an EPHA2 overexpression vector to determine if it could reverse the proliferation impairment caused by METTL1 knockdown. Western Blot results showed that in the METTL1 control group, even with EPHA2 overexpression, the increase in p-AKT expression in the AKT pathway was not as significant compared to when EPHA2 was overexpressed in the METTL1 knockdown background (**Fig. 6A**). A similar trend was observed for pmTOR, although the changes were less pronounced than for p-AKT, indicating that the AKT pathway was reactivated. The CCK-8 proliferation assay showed increased cell proliferation in both the METTL1 control and knockdown groups with EPHA2 overexpression (**Fig. 6B**). The colony formation assay produced similar results (**Fig. 6C**), demonstrating that EPHA2 could rescue the proliferation impairment caused by METTL1 knockdown.Cell cycle analysis further confirmed that EPHA2 overexpression rescued the G0/G1 phase arrest caused by METTL1 knockdown. The METTL1-sh32+EPHA2-NC group had the highest proportion of cells in the G0/G1 phase compared to the control group, with other groups showing intermediate levels (**Fig. 6E**), consistent with the CCK-8 and colony formation assays.

**Figure 6.**
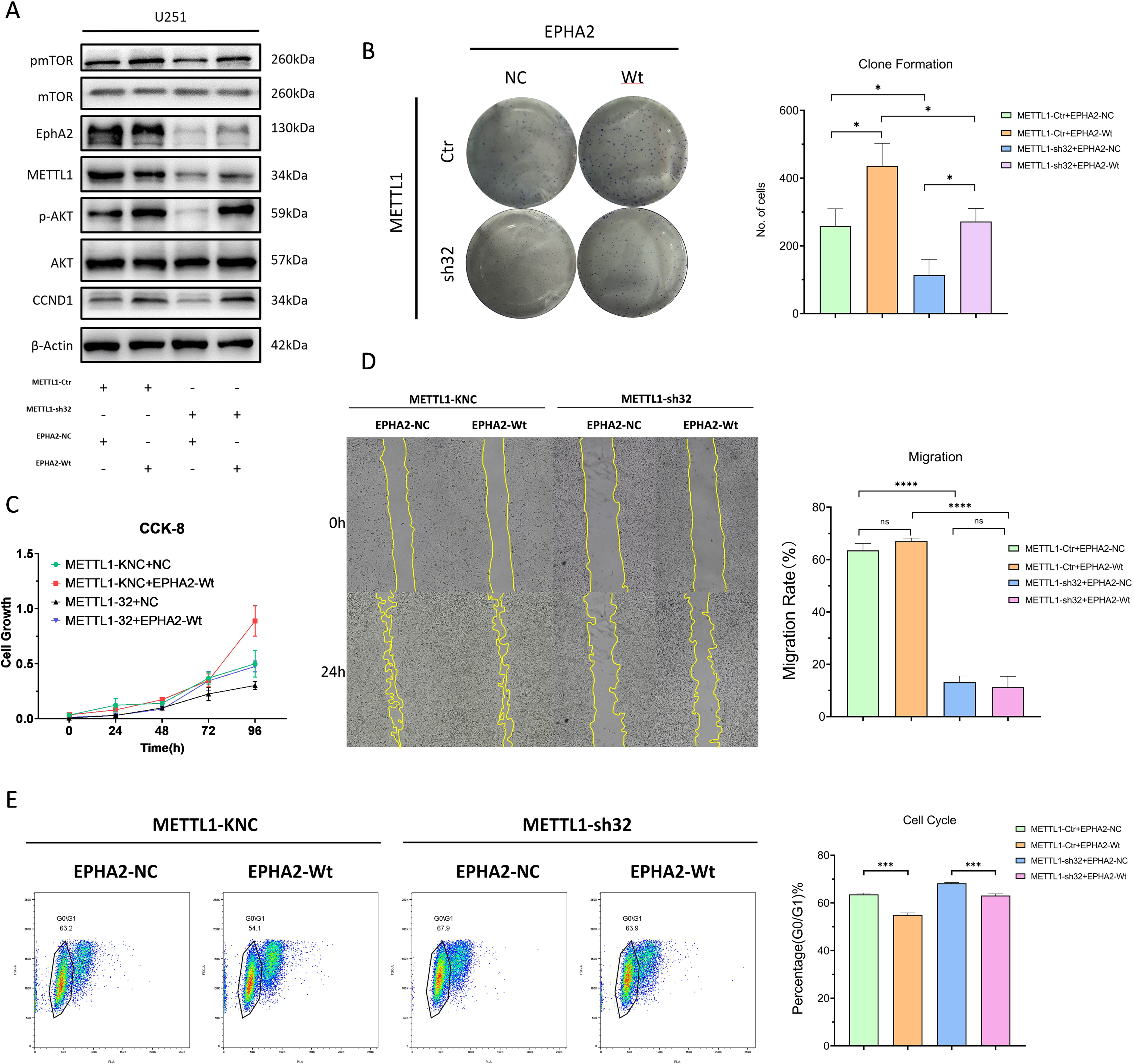
EPHA2 rescue experiment following METTL1 knockdown. A, Western Blot analysis of EPHA2 rescue experiment in METTL1 control and knockdown groups. B, Colony formation assay and quantitative analysis of EPHA2 rescue following METTL1 control and knockdown. C. CCK-8 assay following EPHA2 rescue. D. Scratch assay and quantitative analysis following EPHA2 rescue. E. Cell cycle assay and G0/G1 phase cell population analysis following EPHA2 rescue.

We also explored whether restoring EPHA2 expression could rescue other cell functions like migration. The scratch assay showed that even with EPHA2 overexpression, cell migration remained inhibited (**Fig. 6D**), suggesting that METTL1’s modification of EPHA2 primarily altered cell proliferation, indicating other downstream targets potentially affecting migration and proliferation via m7G modification.

### METTL1 promotes glioma proliferation through EPHA2 in vivo

In vivo experiments supported these findings. Subcutaneous tumors in nude mice derived from METTL1-knockdown cells exhibited slower growth and smaller volumes compared to control groups over 40 days (**Fig. 7A**). Western blot analysis of tumor samples showed reduced expression of PTTG1, EPHA2, p-AKT, p-mTOR, p-ERK1/2, and CCND1 in the METTL1 knockdown group (**Fig. 7A**), implicating these genes and associated pathways in METTL1-mediated glioma progression.

**Figure 7.**
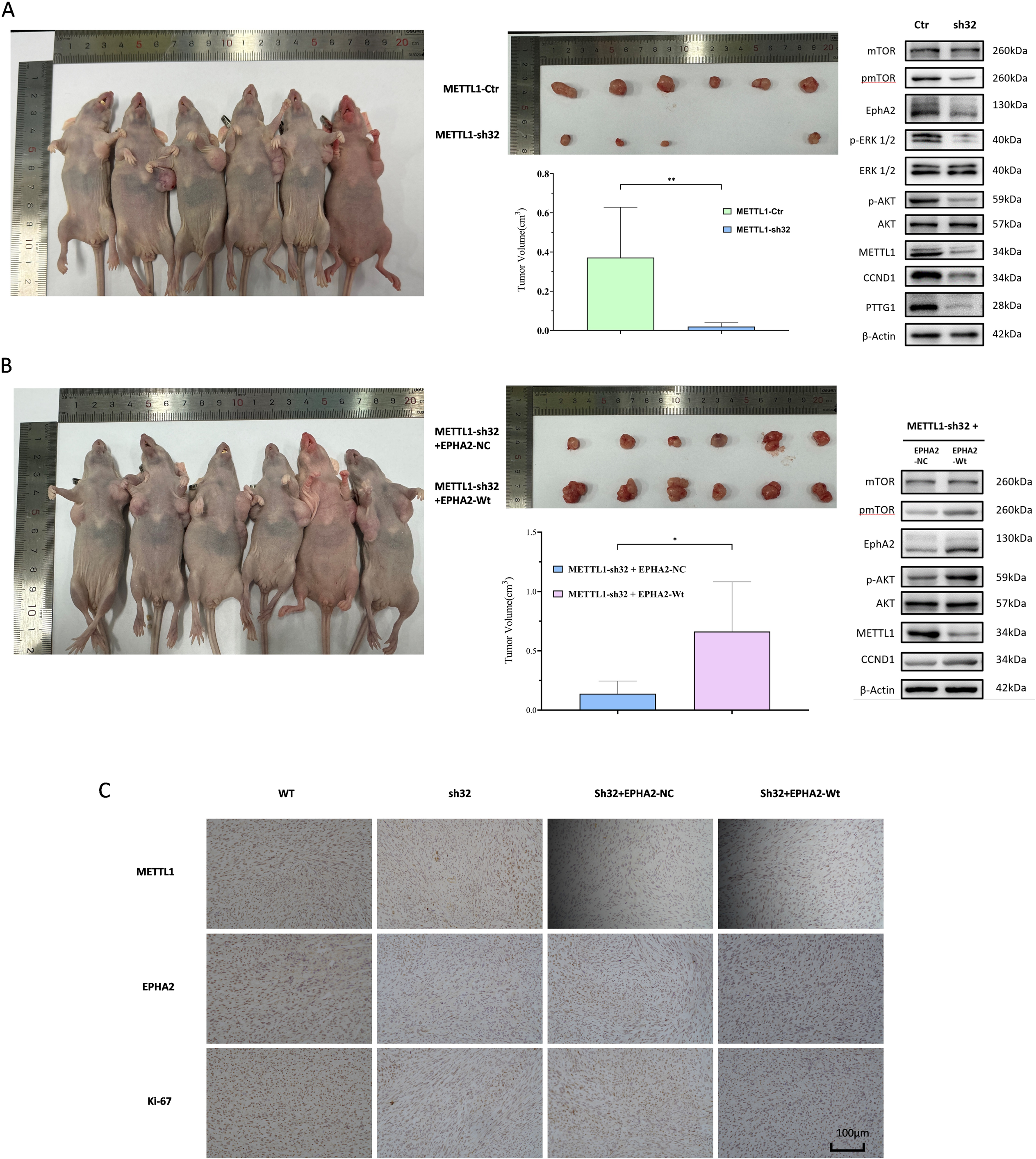
Nude mouse subcutaneous tumor formation rescue experiment. A, Subcutaneous tumor formation in nude mice for the METTL1-Ctr (left) and METTL1-sh32 (right) groups. B, On the left side of the nude mouse, subcutaneous injection of METTL1-sh32 + EPHA2-Wt cells is shown, while on the right side, subcutaneous injection of METTL1-sh32 + EPHA2-NC cells is displayed. C. Representative immunohistochemical staining images of METTL1, EPHA2, and Ki-67 in mouse tumor tissues.

EPHA2 rescue experiments showed that the proliferation impairment due to METTL1 knockdown was partially restored in vivo (**Fig. 7B**). Tumors in the left armpit of mice injected with METTL1-sh32+EPHA2-Wt cells grew faster than those in the right armpit injected with METTL1-sh32+EPHA2-NC control cells. After euthanizing the mice, analysis of the subcutaneous tumors revealed that the tumors in the METTL1-sh32+EPHA2-NC group were smaller compared to those in the METTL1-sh32+EPHA2-Wt group (**Fig. 7B**). Western Blot analysis of subcutaneous tumors indicated that EPHA2 overexpression increased p-AKT and pmTOR levels, reactivating the AKT pathway and restoring Cyclin D1 expression (**Fig. 7B**). Immunohistochemical analysis revealed similar results, with a decrease in Ki-67 expression, a marker of tumor proliferation, following METTL1 knockdown. However, EPHA2 overexpression led to a recovery in Ki-67 expression.

## Discussion

Epigenetics is a complex process involving various modifications, such as histone modifications, chromatin remodeling, nucleosome positioning, DNA methylation, and RNA methylation. Among RNA modifications, over 170 types exist, with methylation being predominant, including m6A, m5C, and m7G.

Current research on the relationship between glioma and METTL1 is limited. Bioinformatics analysis revealed significant enrichment of METTL1 and m7G modifications in gliomas, especially in high-grade glioblastomas, impacting prognosis. This suggests a key role for these epigenetic modifications in glioma development and progression. Western Blot analysis of clinical samples confirmed these findings, prompting further investigation into METTL1’s role in glioma.

We constructed stable METTL1 knockdown and overexpression glioma cell lines to explore METTL1’s impact on glioma progression. Knockdown of METTL1 significantly inhibited cell proliferation, migration, and invasion, with more cells arrested in the G0/G1 phase. Overexpression of METTL1 had less pronounced effects on these phenotypes.

Recent studies indicate that METTL1 adjusts specific RNA translation levels through m7G modification of tRNA, particularly Arg-TCT-4-1(43). While most research focuses on tRNA m7G, m7G sites have been found in mRNA, predominantly in CDS and 3’UTR regions, with fewer in 5’UTR(31). Initial attempts to link METTL1 knockdown effects to tRNA methylation did not align with gene expression changes seen in other models. Notably, METTL1 knockdown also reduced mRNA levels of cell cycle-related genes, highlighting the extensive presence of m7G sites within mRNA. In recent years, with the development of dual-luciferase reporter gene technology, extensive studies have explored the functional roles of mRNA 5’UTR and 3’UTR regions. Particularly in m6A modification research, it has been demonstrated that METTL3 increases mRNA stability through m6A modification in UTR regions, thereby promoting target protein translation, achieving significant progress(44). Drawing on this methodology, similar experimental strategies have been applied in m7G modification studies, leading to preliminary exploration of mRNA UTR region functions. With the recent publication of m7G site sequencing databases for mRNA, it has been observed that some genes have m7G modification sites exclusively in the coding sequence, providing a new direction for further investigation into the functional roles of m7G modification.

EPHA2, a 130kDa transmembrane glycoprotein tyrosine kinase receptor(45), was first discovered in 1990. Under normal conditions, EPHA2 binds to its adjacent ligand ephrin A1, an epidermal growth factor-like molecule expressed on neighboring cells(46), and induces various signaling networks upon cell-cell contact. Unlike EPHA2, ephrin A1 shows reduced expression in several aggressive tumors(47). As a membrane protein, EPHA2 participates in both forward and reverse signaling mediated by its interaction with ephrins, a mechanism referred to as ephrin-EPHA2 bidirectional signaling(48,49). Further studies have revealed that the high expression of EPHA2 in tumor cells exhibits ligand-independent kinase activity due to the lack of regulation by ephrin binding. This partially explains its malignant effects in a non-phosphorylated state(50,51). Additionally, a recent study on patients with histopathologically confirmed primary gliomas revealed strong EPHA2 expression in glioma cells, with its expression negatively correlating with overall survival(52). These findings align with our bioinformatics analysis, further supporting the close association between EPHA2 and gliomas. Using RIP-qPCR and MeRIP-PCR, we validated the presence of credible m7G modification sites on EPHA2 mRNA. RNA stability assays confirmed that METTL1 knockdown reduced the stability of EPHA2 mRNA. The abundance of m7G modification at the EPHA2-k1 site, as detected in MeRIP-qPCR experiments, was lower than at the k2 site. This discrepancy may explain why only one m7G modification site in the CDS region of EPHA2 was identified in the m7G database after data correction.

In our study, although RIP and MeRIP experiments successfully identified m7G modification sites within the CDS region of mRNA, these methods were unable to precisely determine the exact locations of these modifications at the nucleotide level. To further explore and pinpoint these m7G modification sites. We designed synonymous mutations for the predicted methylation-prone G bases within the suspected modification regions, converting them into C bases with similar codon usage frequencies. This approach ensured that the final protein structure remained unaffected while providing a method to examine the specific impact of these G base mutations on m7G modification. We introduced the mutated EPHA2 transcripts into glioma cells and monitored the expression changes of these exogenous EPHA2 transcripts and their mutants under METTL1 knockdown conditions. The results aligned with our expectations, showing that the EPHA2 mRNA with double-site mutations was minimally affected by METTL1 knockdown. Notably, its RNA stability was significantly reduced compared to other transcripts. This finding provides compelling evidence that METTL1 specifically methylates two key sites in the CDS region of EPHA2 mRNA, thereby significantly enhancing its RNA stability and uncovering a critical epigenetic regulatory mechanism.

The regulatory role of EPHA2 in the AKT signaling pathway has been extensively validated across various tumor types, including gliomas(36,45–47). In this study, we utilized METTL1 knockdown and EPHA2 rescue models to investigate the interactions among METTL1, EPHA2, and the AKT pathway. Our results confirmed that METTL1 influences the AKT signaling pathway by regulating EPHA2, thereby affecting glioma cell proliferation.

In addition to EPHA2, PTTG1 and PPIC were identified as potential targets of METTL1-mediated m7G methylation. Although direct point mutation experiments were not conducted to validate their m7G modifications, our observations of RNA modification patterns suggest that the mRNA of PTTG1 and PPIC may play similar regulatory roles through m7G modification. Moreover, m7G modification sites have been identified within the mRNA of many other genes, likely influenced by METTL1-mediated methylation, offering promising directions for future research. This study not only provides new insights into the role of METTL1-mediated m7G modification in RNA stability regulation but also underscores the importance of precisely mapping m7G modification sites to uncover their biological functions.

While we confirmed specific G bases with m7G methylation in exogenous EPHA2 transcripts through synonymous mutations and indirectly compared the expression differences among various transcripts under METTL1 knockdown, this experimental approach provided valuable insights. However, we must acknowledge its limitations. For instance, we cannot entirely exclude the potential influence of tRNA methylation on translation efficiency, which necessitates further validation of the reliability of using the vector’s co-expressed mCherry gene as an exogenous reference. Additionally, METTL1 may possess other unexplored functions in gliomas, presenting new directions for investigation. The observed changes in glioma cell migration and invasion abilities under METTL1 knockdown highlight the need to explore the underlying mechanisms of these phenomena. Furthermore, understanding how METTL1-mediated m7G methylation in mRNA CDS regions affects mRNA stability remains an unresolved question, warranting deeper mechanistic studies.

## Conclusion

This study found that METTL1 is highly expressed in gliomas and is positively correlated with poor prognosis in patients. High METTL1 expression promotes key phenotypes such as cell proliferation, migration, and invasion. Additionally, we discovered that EPHA2 is regulated by METTL1 through m7G methylation in the CDS region. This m7G modification increases the stability of EPHA2 mRNA, subsequently affecting the AKT signaling pathway and enhancing the proliferative capacity of glioma cells.

## List of abbreviations

METTL1: Methyltransferase-like 1
GTEx: The Genotype-Tissue Expression
CCLE: Cancer Cell Line Encyclopedia
TCGA: The Cancer Genome Atlas
CGGA: Chinese Glioma Genome Atlas
CNS: Central Nervous System
GO: Gene Ontology
GSEA: Gene Set Enrichment Analysis
KEGG: Kyoto Encyclopedia of Genes and Genomes
IDH: Isocitrate Dehydrogenase
AUC: Area Under the Curve
WHO: World Health Organization
IHC: Immunohistochemistry
C-index: Consistency Index
qPCR: Quantitative Real-time PCR
DMEM: Dulbecco’s Modified Eagle Medium
FBS: Fetal Bovine Serum
PBS: Phosphate-buffered Saline
PVDF: Polyvinylidene Fluoride
PAGE: Polyacrylamide Gel Electrophoresis
LGG: Low-grade Gliomas
GBM: Glioblastoma Multiforme
OS: Overall Survival
DSS: Disease-specific Survival
KD: Knockdown
OE: Overexpression
WB: Western Blot
siRNA: Small Interfering RNA
AKT: Protein Kinase B
PTTG1: Pituitary Tumor-Transforming 1
EPHA2: Erythropoietin-producing Hepatoma Receptor A2
MeRIP-PCR: Methylated RNA Immunoprecipitation-PCR
RIP: RNA Immunoprecipitation
shRNA: Short Hairpin RNA
CCK8: Cell Counting Kit-8
DMSO: Dimethyl Sulfoxide.

## Declarations

### Ethics approval and consent to participate

This study was approved by the Education and Ethics Committee of the First Affiliated Hospital of China Medical University. Informed consent was obtained from all participants. All animal experimental procedures were conducted in accordance with the National Institutes of Health Guide for the Care and Use of Laboratory Animals, and were approved by the Animal Ethics Committee of China Medical University.

## Consent for publication

Not applicable.

## Availability of data and material

The datasets generated and/or analyzed during the current study are available in UCSC Xena (http://xena.ucsc.edu/) and CGGA (http://www.cgga.org.cn).

## Authors’ contributions

TTX and GYL were responsible for the experimental design, execution, and manuscript writing. PY, YQS, provided technical support for the experiments. XF, JHH, CRX, JHL and SXY provided bioinformatics analysis support. All authors approved the final manuscript.

## Acknowledgements

Not applicable.

---

Supplementary Fig. S1 A. Construction of METTL1 knockdown stable cell lines in glioma cell lines and evaluation of infection efficiency. B. Statistical analysis of colony formation assay. C. qPCR was used to assess the efficiency of METTL1 knockdown and overexpression in U251 and LN229 cell lines. D. In stably transfected U251 and LN229 cells, compared to the EV group, METTL1 overexpression in the OE group did not significantly affect cell migration ability. E. Bar chart showing the statistical analysis of the number of transmembrane cells in the Transwell assay (left) and Invasion assay (right).

Supplementary Fig. S2 A. Survival differences of EPHA2, PTTG1, and PPIC with different expression levels in glioma patients from the TCGA and CGGA databases, with higher METTL1 expression associated with poorer prognosis. B. Differential expression analysis of the three genes in LGG, GBM, and normal brain tissues, showing the highest expression levels in the GBM group, followed by the LGG group. C. RNA correlation analysis between METTL1 and EPHA2, PTTG1, and PPIC. D-E. PTTG1 and PPIC were identified as potential METTL1 modification targets, with a single possible modification site located in the CDS region.

Supplementary Fig. S3 A. Knockdown of METTL1 in U251 cells led to a decrease in the mRNA expression levels of PTTG1 and PPIC. B. RIP assay in U251 and LN229 cell lines showed that the mRNA abundance pulled down with the METTL1 antibody was higher compared to IgG. C. Primers designed for detection of mCherry on the lentiviral vector.

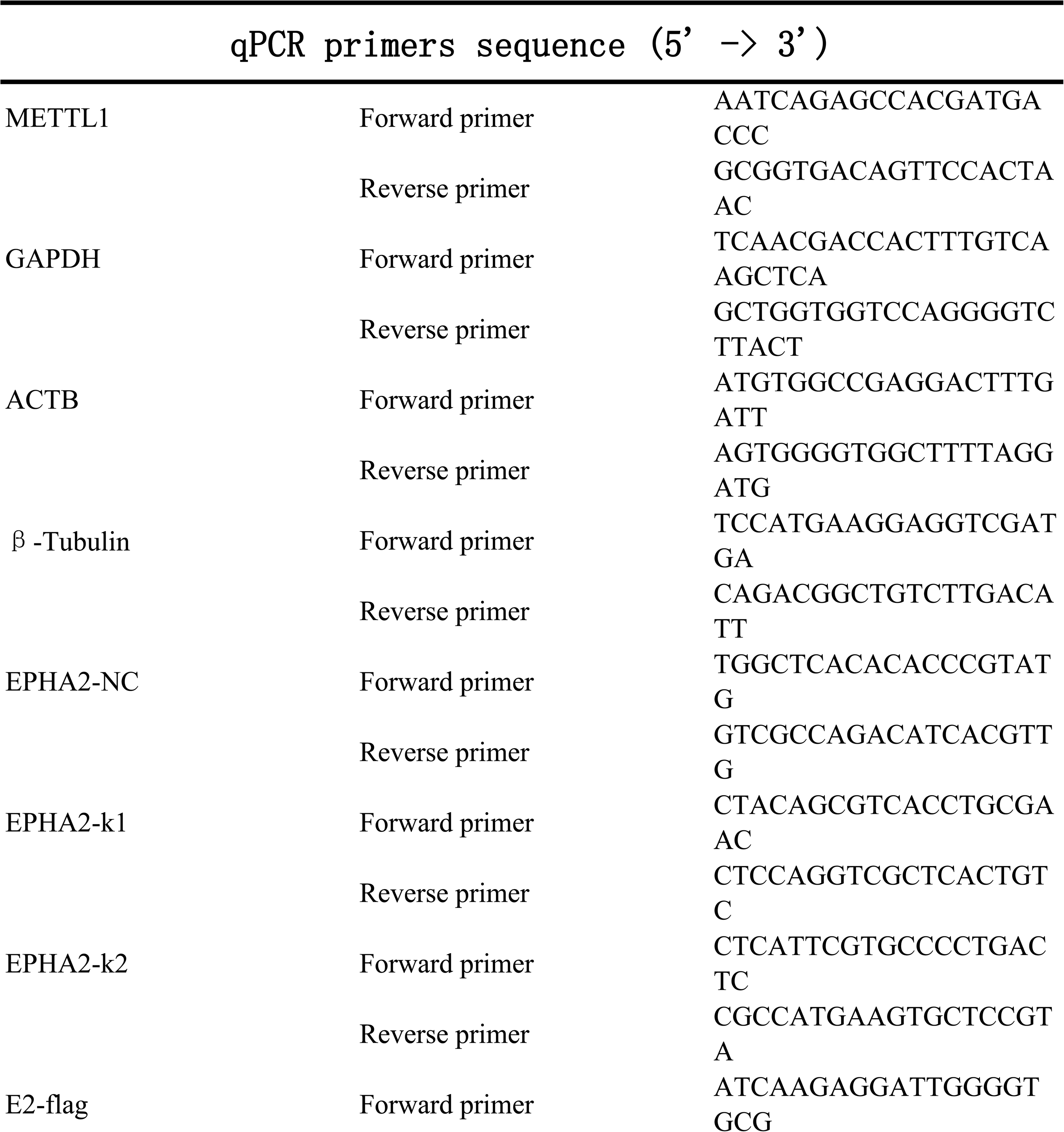
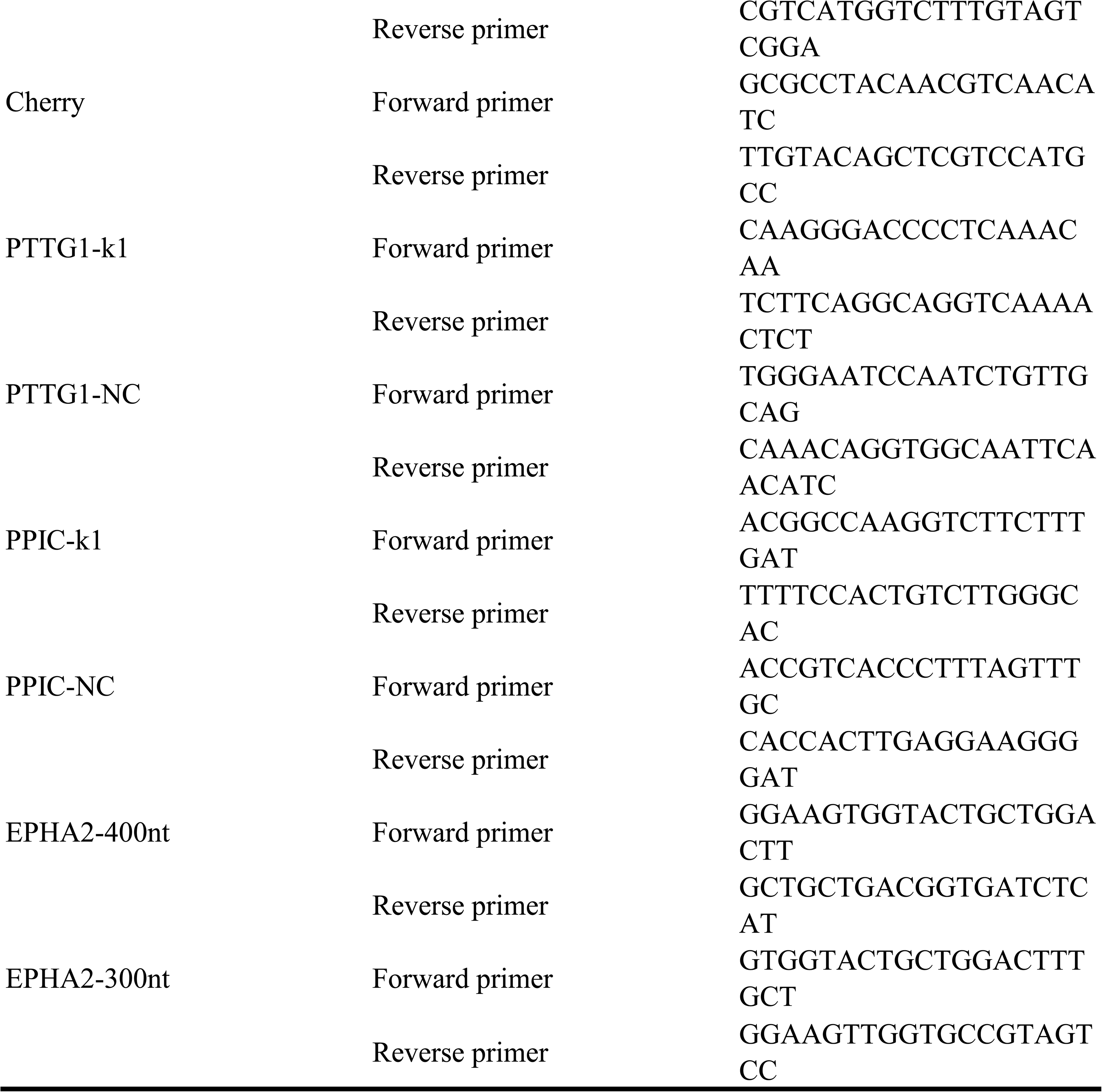

